# Loss of mCHH islands in maize chromomethylase and DDM1-type nucleosome remodeler mutants

**DOI:** 10.1101/253567

**Authors:** Fang-Fang Fu, R. Kelly Dawe, Jonathan I. Gent

## Abstract

Plants make use of three types of DNA methylation, each characterized by distinct DNA methyltransferases. One type, RNA-directed DNA methylation (RdDM), is guided by siRNAs to the edges of transposons that are close to genes, areas called mCHH islands in maize. Another type, chromomethylation, is guided by histone H3 lysine 9 methylation to heterochromatin across the genome. We examined DNA methylation and small RNA expression in plant tissues that were mutant for both copies of the genes encoding chromomethylases as well as mutants for both copies of the genes encoding DDM1-type nucleosome remodelers, which facilitate chromomethylation. Both sets of double mutants were nonviable but produced embryos and endosperm. RdDM was severely compromised in the double mutant embryos, both in terms of DNA methylation and siRNAs. Loss of 24nt siRNA from mCHH islands was coupled with a gain of 21, 22, and 24nt siRNAs in heterochromatin. These results reveal a requirement for both chromomethylation and DDM1-type nucleosome remodeling for RdDM in mCHH islands, which we hypothesize is due to dilution of RdDM components across the genome when heterochromatin is compromised.

## INTRODUCTION

### RdDM and other forms of DNA methylation in plants

Chromatin modification directed by RNA interference (RNAi) and related processes is essential to genome defense in most eukaryotes. The molecular mechanisms vary even within the same cell, but key features are short interfering RNAs (siRNAs) which guide argonaute (Ago) proteins and induce methylation of histone H3 lysine 9 (H3K9me) (Holoch and Moazed, 2015). RNA-directed DNA methylation (RdDM) is a well-characterized example in plants (Cuerda-Gil and Slotkin, 2016). RdDM leads to H3K9me2 (Jackel et al., 2016; Fultz and Slotkin, 2017); but as its name suggests, is better known for methylating DNA. At least two *Arabidopsis thaliana* proteins that function in RdDM physically interact with a DNA methyltransferase, indicating a direct connection between RdDM and DNA methylation (Gao et al., 2010; Zhong et al., 2014). RdDM represses repetitive and foreign DNA but can also influence gene expression, at gene regulatory elements for example (Rowley et al., 2017), and is involved in epigenetic phenomena such as paramutation (Hollick, 2017) and genomic imprinting, (Satyaki and Gehring, 2017).

DNA methylation occurs at cytosines in all sequence contexts and is catalyzed by three distinct types of methyltransferases in plants (Du et al., 2015). The first, related to Arabidopsis METHYLTRANSFERASE1 (MET1), which is homologous to mammalian DNMT1, is responsible for replication-coupled CG methylation. The second, related to Arabidopsis CHROMOMETHYLASE (CMT1, 2, and 3), methylate CHGs and CHHs, where H is A, T, or C. The chromodomains of these methyltransferases guide their activity to regions of H3K9me1 or H3K9me2. The third, related to Arabidopsis DOMAINS REARRANGED METHYLTRANSFERASE (DRM1 and 2), methylates cytosines in all sequence contexts through RdDM. Depending on the species, methylation in the CHH context (mCHH) can be used as an indicator of RdDM, as mCHH produced by CMT is of lesser magnitude than that of RdDM (Niederhuth et al., 2016). Division of methylation contexts into mCG, mCHG, and mCHH is a helpful simplification, but methyltransferases have additional nucleotide preferences (Gouil and Baulcombe, 2016).

### Repressive chromatin modifications promote RdDM

Although RdDM was discovered as a means to initiate methylation at previously unmethylated DNA (referred to as de novo methylation (Wassenegger et al., 1994)), discoveries since then have revealed that most RdDM activity occurs at already-methylated and repressed loci (Cuerda-Gil and Slotkin, 2016). This maintenance form of RdDM, called canonical RdDM, depends on the activity of the RNA polymerase II variants Pol IV and Pol V and is responsible for the majority of 24nt siRNAs and resulting DNA methylation. Canonical RdDM in Arabidopsis requires methylated histone H3 lysine 9 (H3K9me) and unmethylated lysine 4 (H3K4) to be fully effective (Johnson et al., 2008; Kuhlmann and Mette, 2012; Greenberg et al., 2013; Law et al., 2013; Zhang et al., 2013; Johnson et al., 2014; Stroud et al., 2014; Li et al., 2015b). Histone deacetylation is also required (Blevins et al., 2014). DNA methylation also promotes RdDM, as *drm2* and *cmt3* mutants have reduced levels of 24nt siRNAs (Law et al., 2013; Stroud et al., 2014; Li et al., 2015b). Loss of H3K9me2 in these mutants may explain the loss of RdDM, as H3K9me2 recruits Pol IV (Law et al., 2013; Zhang et al., 2013). DNA methylation may also promote RdDM independently of H3K9me2, however, as other RdDM components link DNA methylation to Pol V (Johnson et al., 2014; Liu et al., 2014).

### Heterochromatin inhibits RdDM

RdDM, by definition, relies on transcription, so the fact that histone modifications that inhibit transcription promote RdDM is surprising. Repetitive DNA elements in the genome is repressed in multiple different genomic contexts, e.g., large regions of heterochromatin, heterochromatin-euchromatin boundaries, or small regions of repressive chromatin in larger euchromatic contexts (Sigman and Slotkin, 2016). The heterochromatic middle regions of long transposons are depleted of RdDM relative to their euchromatin-flanking ends, which tend to be enriched for RdDM (Lee et al., 2012; Zhong et al., 2012; Zemach et al., 2013; Stroud et al., 2014; Li et al., 2015b). The enrichment for RdDM at heterochromatin-euchromatin boundaries is especially clear in maize because its heterochromatin and euchromatin are highly interspersed (Gent et al., 2013; Gent et al., 2014; Li et al., 2015a; Niederhuth et al., 2016).

That heterochromatin inhibits RdDM is suggested not only by its distribution in the genome, but also by activation of RdDM in normally heterochromatic regions in plants that lack the SNF2 family nucleosome remodeling protein DECREASED DNA METHYLATION1 (DDM1) (Nuthikattu et al., 2013; Creasey et al., 2014; McCue et al., 2015; Panda et al., 2016), known as LYMPHOID SPECIFIC HELICASE (LSH) in mammals (Dennis et al., 2001). DDM1 is required for access of MET1 and CMT-type methyltransferases to non-transcribed, nucleosome-bound DNA in Arabidopsis (Lyons and Zilberman, 2017). The vegetative cell of pollen in Arabidopsis provides another line of evidence that heterochromatin inhibits RdDM because this cell type undergoes a dramatic decondensation of heterochromatin and activation of RdDM (Schoft et al., 2009; Slotkin et al., 2009; Mérai et al., 2014). DDM1 may also be reduced or absent from vegetative cells, as transgene driven expression of a DDM1 fusion protein from a *ddm1* promoter produced signal in sperm but not in vegetative cells (Slotkin et al., 2009). Vegetative cells have increased mCHH (Calarco et al., 2012; primarily driven by CMT2, but also DRM2 activity (Hsieh et al., 2016).

The apparent contradiction between chromatin modifications associated with heterochromatin promoting RdDM, yet heterochromatin itself inhibiting RdDM may be explained by the relative abundance of such modifications: Moderate levels of H3K9me2, for example, may promote RdDM, while dense H3K9me2 may block transcription. Additional modifications or higher-order chromatin structure that affects chromatin accessibility to RNA Pol V and Pol IV may also contribute.

### Relationship between RdDM and heterochromatin in maize

Its quick life cycle, small genome, and resilience to loss of DNA methylation have made Arabidopsis the plant model of choice for research on DNA methylation and chromatin. The discoveries made with Arabidopsis have been tremendously helpful in understanding similar phenomena in more difficult to work with plants such as maize, with its large, repetitive genome and nonviable methylation mutants (Li et al., 2015a). However, the differences between maize and Arabidopsis also limit the extent to which results can be projected from Arabidopsis to maize. Maize siRNA size distributions are different from Arabidopsis in the abundance of 22nt siRNAs (Nobuta et al., 2008; Wang et al., 2009), and the presence of abundant meiotic phasiRNAs (Johnson et al., 2009; Zhai et al., 2015). Maize lacks a CMT2-type chromomethylase (Zemach et al., 2013; Bewick et al., 2017). It has multiple copies of the major subunits of Pol IV and Pol V complexes with potential for specialized functions (Haag et al., 2014).

Here, we endeavored to determine the relationship between RdDM and other forms of DNA methylation using DNA methylation mutants in maize. In particular, we focused on DDM1 and the chromomethylases given their major effects on heterochromatin in Arabidopsis (Gendrel et al., 2002). The maize genome encodes two chromomethylases, named ZMET2 and ZMET5 (also known as DMT102 and DMT105) (Li et al., 2014). Both are functionally more similar to CMT3 than to CMT1 or CMT2 (Bewick et al., 2017). The maize genome also encodes two DDM1-like nucleosome remodelers, CHR101 and CHR106 (Li et al., 2014). The effects of single mutants of all four genes on whole genome methylation have been investigated previously (Gent et al., 2014; Li et al., 2014; Gouil and Baulcombe, 2016). Single mutants of *chr101* and *chr106* have little effect on DNA methylation, while single mutants of *zmet2* and *zmet5* have decreased mCHG in both leaves and developing ears, and decreased mCHH in leaves but not in developing ears. Double mutants of *chr101* and *chr106* and double mutants of *zmet2* and *zmet5* are nonviable (Li et al., 2014). We found that double mutants can produce embryos and endosperm, however, and we examined DNA methylation and siRNAs in both tissues. RdDM was severely compromised in developing embryos, with near complete loss of both 24nt siRNAs and mCHH from mCHH islands. The loss of 24nt siRNAs from mCHH islands was accompanied by dramatic gains of 21 and 22nt siRNAs at heterochromatic loci in the genome but these siRNAs did not direct DNA methylation.

## RESULTS

### Generation of *ddm1* double and *cmt* double mutant embryo and endosperm

To make plants that lacked chromomethylases or DDM1-type nucleosome remodelers, we obtained UniformMu stocks with *Mu* insertions in exons of *Zmet2* (*mu1013094*) (Gent et al., 2014) and *Zmet5* (*mu1017456*), and in the DDM1 genes *Chr101* (*mu1044815*) and *Chr106* (*mu1021319*). Here we will refer to *zmet2 zmet5* homozygous double mutants as *cmt,* and the *chr101* and *chr106* double mutants as *ddm1*. Mutants that carried a single wildtype copy of either *Zmet2* or *Zmet5* were viable and fertile, and crosses between such mutants produced kernels with sectors of pigmented aleurone (Supplemental Figure 1). This phenotype is characteristic of *Mutator* transposon activity in UniformMu stocks, where excision of a *Mu* from the *bz1-mum9* allele of the *Bz1* gene can restore pigmentation in small sectors during development (McCarty et al., 2005). *Mu* activation has been observed in a maize mutant lacking the RdDM component MEDIATOR OF PARAMUTATION 1 (MOP1) (Woodhouse et al., 2006). We found a tight correlation between pigmented sectors in the aleurone and *zmet2 zmet5* (*cmt*) double mutant genotype (Supplemental Table 1). The *cmt* kernels contained both endosperm and embryos but were usually incapable of more than a couple centimeters of root development upon germination. A second pair of *zmet2* and *zmet5* alleles (*zmet2-m1* and *zmet5-m1)* that was introgressed into the B73 genetic background produced homozygous double mutant kernels at the expected ratios and were also nonviable (Supplemental Table 2). Failure to produce double mutant plants with these alleles was previously reported (Li et al., 2014)). These kernels lacked the *Mu* insertion in the *Bz1* gene, so were incapable of pigmentation sectoring. Mutants that carried a single wildtype copy of either *Chr101* or *Chr106* were viable and fertile but produced homozygous double mutant (*ddm1*) kernels that were nonviable and had small embryos (Supplemental Table 1 and Supplemental Figure S1). Although in the *bz1-mum9* background, these *ddm1* kernels did not exhibit the sectoring phenotype of *cmt* mutants. They also did not germinate, not even to produce a root tip. A second set of *chr101* and *chr106* alleles (*chr101-m3* and *chr106-m1*) that was introgressed into B73 did not produce any homozygous double mutant kernels (Supplemental Table 2), consistent with the prior study (Li et al., 2014).

### Loss of DNA methylation in mCHH islands in *cmt* and *ddm1* embryo and endosperm

To determine the effects of the mutations on DNA methylation in mature endosperm, we carried out whole genome bisulfite sequencing (WGBS) using the methylC-seq method (Urich et al., 2015) in three tissues: mature endosperm, developing endosperm 14 days after pollination (14-DAP endosperm), and 14-DAP embryos. In *cmt* mutants, mCHG was reduced to near background levels near genes in all three tissues, with little or no effect on mCG (Figure 1). In *ddm1* both mCHG and mCG were mildly reduced (40% reduction in mCG, 50% in mCHG in 14-DAP embryos). These effects on mCHG (nearly absent in *cmt*) and mCHG and mCG (reduced in *ddm1*) are consistent with the expected roles of chromomethylases and DDM1-like nucleosome remodelers in transcriptionally silent heterochromatin. Unexpectedly, we found that RdDM was nearly absent in *cmt* 14-DAP embryos, strongly reduced in mature endosperm, and slightly reduced in 14-DAP endosperm, as evidenced by loss of mCHH (Figure 1). The effect on mCHH could not be explained by background mutations in the UniformMu-derived *cmt* mutant, as 14-DAP sibling embryos with a single wildtype copy of either *Zmet2* or *Zmet5* had near wildtype levels of mCHH (Supplemental Figure S2). RdDM was also nearly absent in *ddm1* 14-DAP embryos, and strongly reduced in both 14-DAP and mature endosperm (Figure 1). The effect is more pronounced when considered in the context of the fact that ZMET2 and ZMET5 can methylate CHH, particularly in the CAA and CTA contexts (Gouil and Baulcombe, 2016). The mCCH subset of mCHH is more specific to RdDM. The mCCH profile in *ddm1* clearly indicated loss of RdDM (Figure 1). This loss of RdDM was also unlikely to be explained by background mutations in the UniformMu-derived *ddm1* mutant, as methylation levels in mature endosperm were high when a single wildtype copy of either *Chr101* or *Chr106* was present (Supplemental Figure S3).

**Figure 1:**
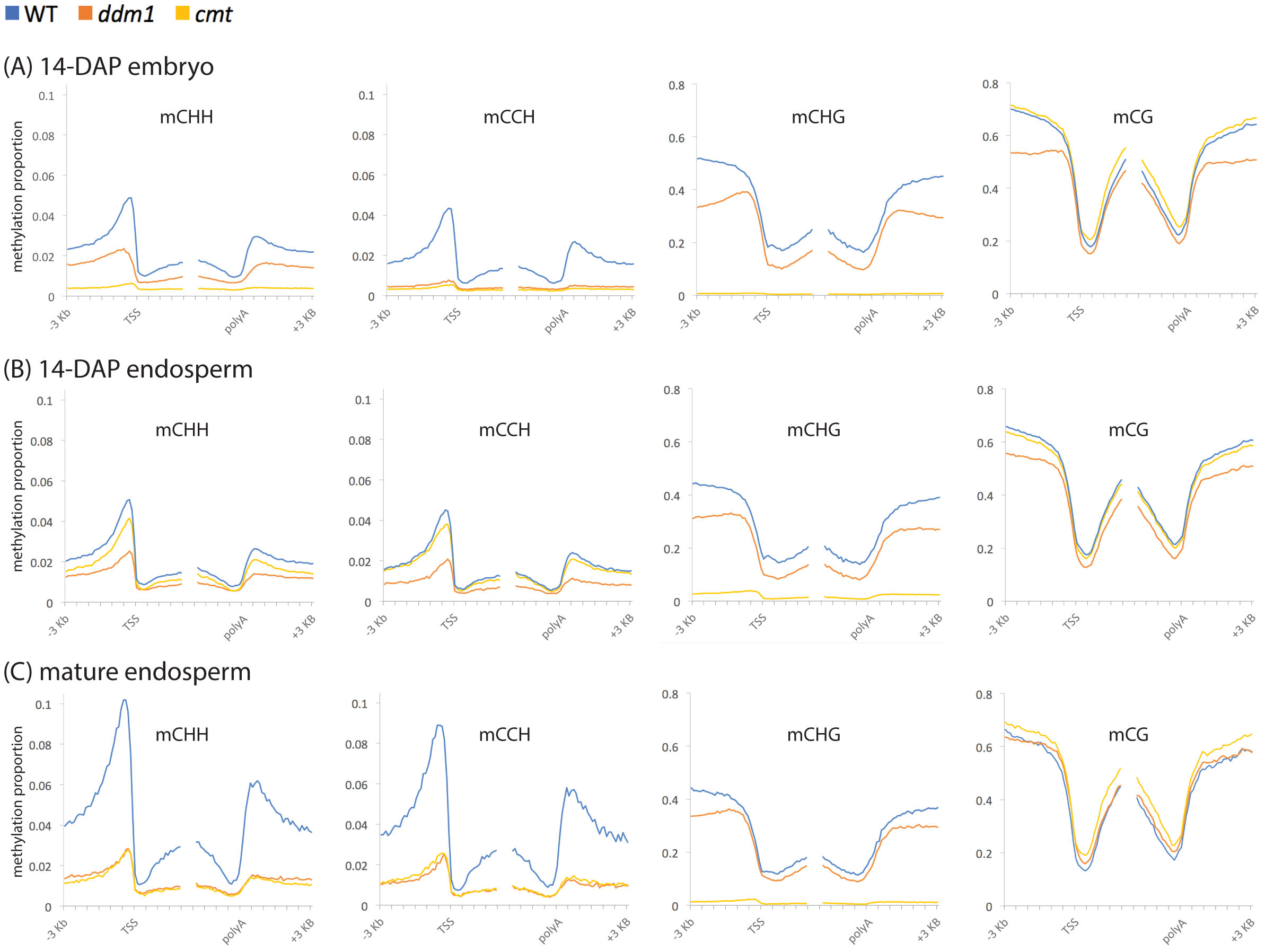
DNA methylation profiles near genes in double mutants **(A)** Methylation in 14-DAP embryo. All genes were defined by their annotated transcription start sites (TSS) and polyadenylation sites (polyA) and split into nonoverlapping lOObp intervals. Methylation for each sample was calculated as the proportion of methylated C over total C in each sequence context (CHH, CCH, CHG, and CG) averaged for each lOObp interval. **(B)** Methylation in 14-DAP endosperm, as in (A) **(C)** Methylation in mature endosperm, as in (A)

### Loss of 24nt siRNAs in mCHH islands in *cmt* and *ddm1* developing embryo and endosperm

Since 24nt siRNAs direct DNA methylation, we sequenced small RNA from 14-DAP endosperm and embryo. To quantify siRNA abundance, we normalized siRNA counts by microRNA (miRNA) counts. We included all mappable small RNAs, both uniquely-mapping and multi-mapping. There was a nearly complete loss of 24nt siRNAs from gene flanks in both *cmt* and *ddm1* 14-DAP embryos (Figure 2A). The loss of 24nt siRNAs was stronger in *ddm1* than in *cmt* 14-DAP endosperm, consistent with the distribution of mCHH and mCCH (Figure 1). A loss of 24nt siRNAs was also evident from the distribution of total siRNA lengths: In homozygous wildtype individuals and heterozygous mutants, the dominant siRNA length was 24nt, but in *cmt* and *ddm1* it shifted to 22nt and to a lesser extent 21nt (Figure 2B). DNA transposons with terminal inverted repeats (TIRs) of the *Harbinger*, *Mutator*, *hAT*, and *Mariner* superfamilies are enriched in mCHH islands (Gent et al., 2013). 24nt siRNAs from these TIR transposons were reduced about 8-fold in *cmt* and in *ddm1* (Figure 2B).

**Figure 2:**
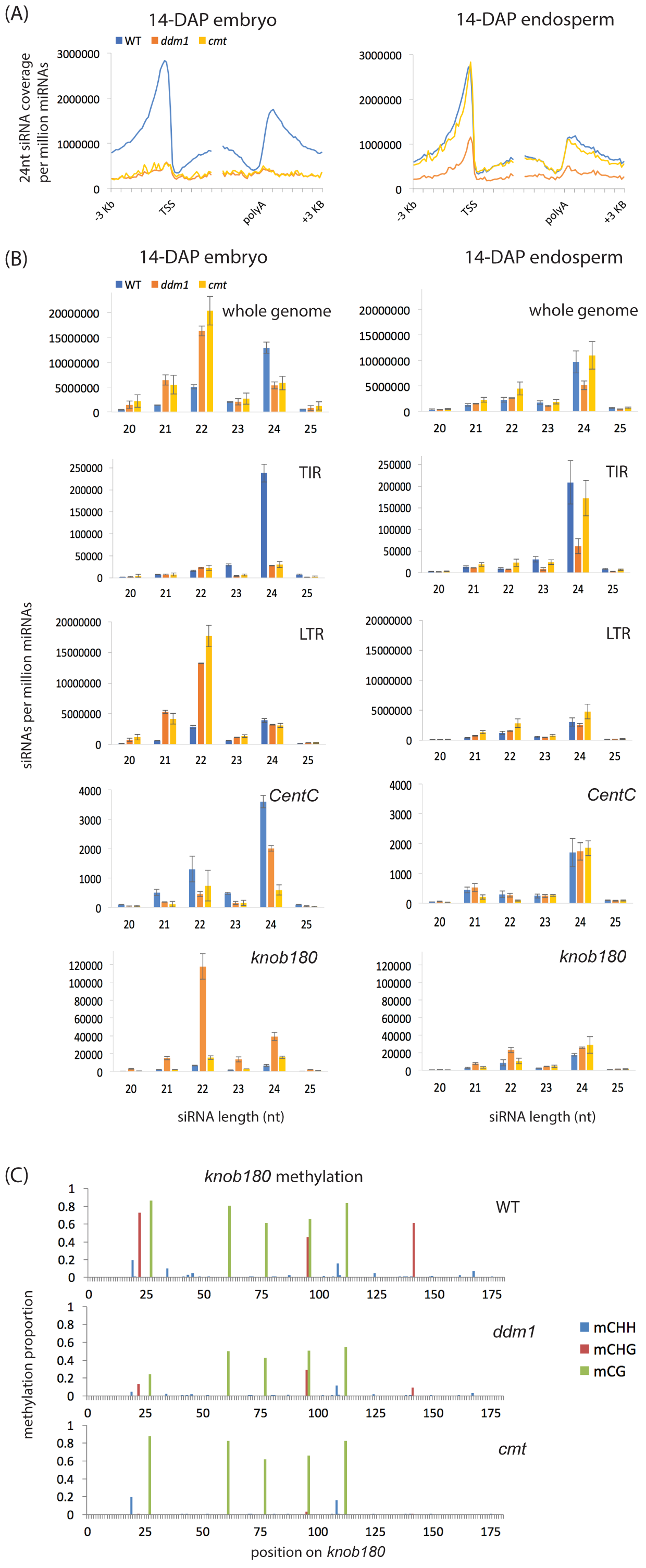
Loss of 24nt siRNAs and gain of 21nt and 22nt siRNAs in double mutants **(A)** 24nt siRNA coverage near genes. All genes were defined by their annotated transcription start sites (TSS) and polyadenylation sites (polyA) and split into nonoverlapping lOObp intervals. The siRNA coverage for each lOObp interval was summed for the complete set of genes and normalized per million miRNAs. **(B)** siRNA lengths, “whole genome” includes all mapped siRNAs. "TIR" includes all siRNAs that overlapped by at least half their lengths to *Mutator, hAT, Harbinger,* or *Mariner* terminal inverted repeat transposons. “LTR” includes all siRNAs that overlapped by at least half their lengths with *Gypsy* or *Copia* long terminal repeat (LTR) retrotransposons. *CentC* includes all siRNAs that aligned to a *CentC* consensus sequence. *knob!80* includes all siRNAs that aligned to a *knob180* consensus sequence. siRNA counts were normalized per million miRNAs. Error bars are standard errors of the means for the biological replicates of each genotype. **(C)** Single-bp DNA methylation in *knob180* repeats. WGBS reads were mapped to the *knob180* consensus sequence, and methylation calculated as the proportion of methylated C over total C in each sequence context.

### Gain of siRNAs in heterochromatin in *cmt* and *ddm1*

Retrotransposons with long terminal repeats (LTRs), which are relatively depleted in mCHH islands, gained both 21nt and 22nt siRNAs in the mutants (8.8 fold gain of 21nt siRNAs in *ddm1*, 7.0 fold gain of 21nt siRNAs in *cmt*; 4.6 fold gain of 22nt siRNAs in *ddm1*, 6.2 fold gain of 22nt siRNAs in *cmt* (Figure 2B)). We also examined siRNAs at two types of high copy tandem repeats: centromeric *CentC* and non-centromeric *knob180*. Both types are depleted of mCHH and siRNAs in wildtype conditions (Gent et al., 2012; Gent et al., 2014), but in the *ddm1* 14-DAP embryos, *knob180* produced 8-fold more 21nt siRNAs, 18-fold more 22nt siRNAs, and 6-fold more 24nt siRNAs than in wildtype (Figure 2B). This increase in *knob180* siRNAs was not accompanied by an increase in mCHH (Figure 2C). A chromosome-level view of 24nt-siRNA coverage showed enrichment towards chromosome arms in wildtype 14-DAP endosperm and embryo, corresponding to gene density and mCHH islands, whereas 21 and 22nt siRNAs had a more uneven distribution with large numbers of siRNAs at discrete loci (Figure 3A). In *cmt* and *ddm1* 14-DAP embryos, 24nt siRNAs showed a distribution similar to wildtype 21 and 22nt siRNAs, with a strong reduction at the majority of loci across the genome. We split the genome into 100bp bins and counted the number of 24nt siRNAs that mapped to each bin (requiring at least 14 bp of the siRNA overlap with the bin). Any locus with at least 5 overlapping siRNAs per 500000 miRNAs, and with siRNAs spanning at least 50 of the 100 bp, was defined as a 24nt siRNA locus. In wildtype 14-DAP embryos, 176342 loci met these criteria, while only 26519 did in *ddm1* embryos and 26546 did in *cmt* embryos. We also identified the subset of 17985 novel *ddm1* 24nt siRNAs that did not meet the criteria in wildtype. Despite the *ddm1* and *cmt* 24nt siRNA loci being defined solely by 24nt siRNAs, they were more strongly enriched for 21 and 22nt siRNAs than 24nt siRNAs (Figure 3B). Similar to the *knob180* tandem repeat (Figure 2C), the *cmt*, *ddm1,* and novel 24nt siRNA loci were highly methylated in mCG and mCHG and poorly methylated in mCHH in wildtype, and did not gain mCHH in either mutant (Figure 3C).

**Figure 3:**
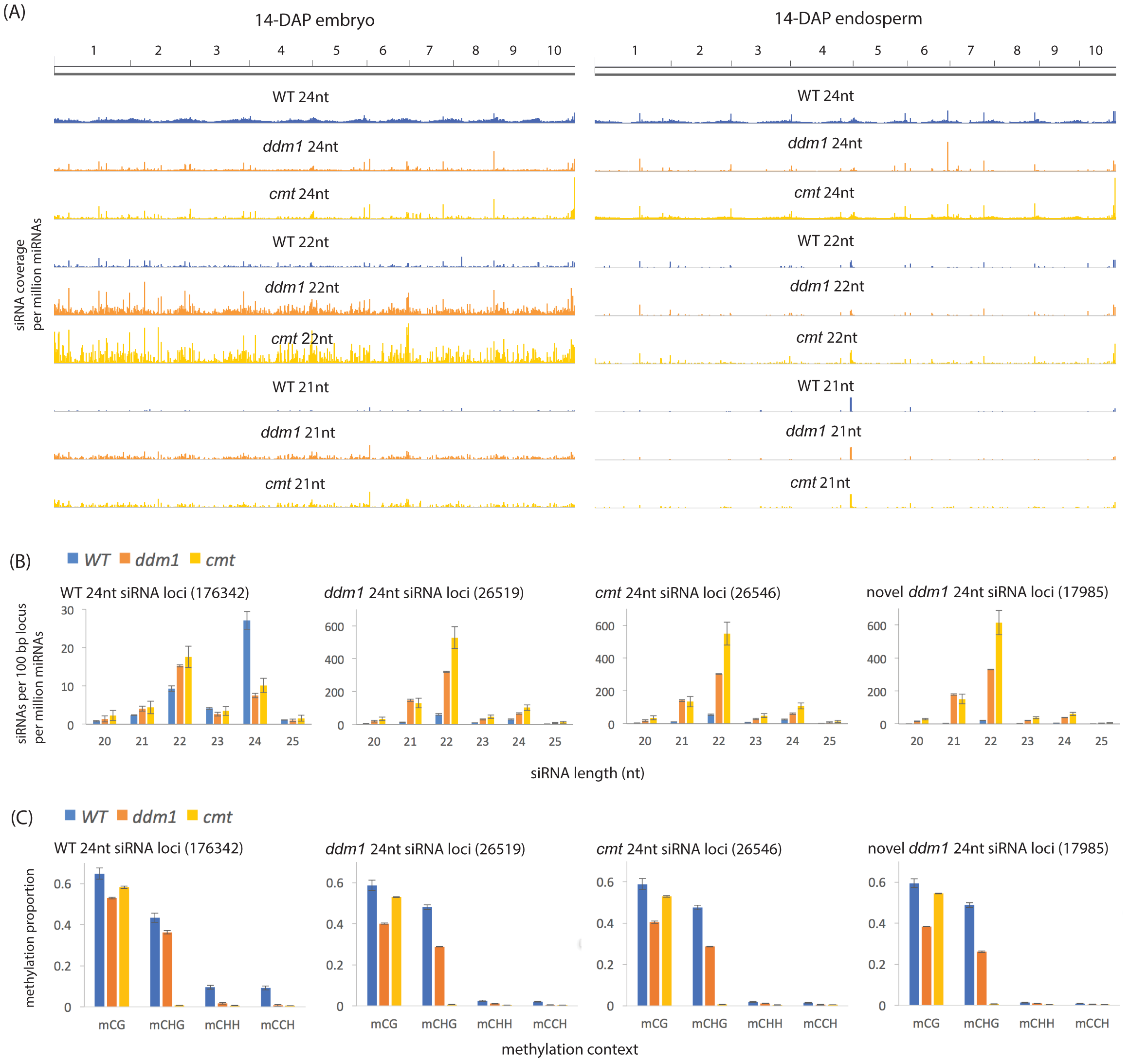
24nt siRNA loci in mutants **(A)** Whole-chromosome siRNA patterns. siRNA coverage counton 500Kb intervals (normalized by miRNA count) is shown for each of the ten chromosomes for 14-DAP embryo and 14-DAP endosperm. Coverage is on the same scale in every track. **(B)** siRNA lengths in 24nt siRNA loci. siRNA counts were normalized by the number of loci in each set (shown in parentheses). All siRNAs that overlapped by at least half their lengths with each type of locus were included. Error bars are standard errors of the means for the biological replicates of each genotype. **(C)** DNA methylation in 24nt siRNA loci. Average methylation for each set of loci was calculated as the proportion of methylated C over total C in each sequence context. Error bars are standard errors of the means for the biological replicates of each genotype.

## DISCUSSION

We found that double mutants of the chromomethylases *zmet2* and *zmet5* (*cmt*) and double mutants of the DDM1-type nucleosome remodelers *chr101* and *chr106 (ddm1*) were deficient in canonical RdDM, as indicated by loss of DNA methylation and 24nt-siRNAs in mCHH islands. The loss of mCHG in *cmt* indicates that *mu1013094* and *mu1017456* are null alleles. The smaller loss of mCHG/mCG in *ddm1* is consistent with null alleles of *ddm1* in Arabidopsis and rice (Zemach et al., 2013; Tan et al., 2016; Lyons and Zilberman, 2017), but in the absence of a clear expectation for a *ddm1* phenotype in maize, residual DDM1 activity in *mu1044815* or *mu1021319* is theoretically possible. The complete nonviability of the *ddm1* kernels indicates that the mutants are at least severe hypomorphs.

In *cmt* 14-DAP embryos, methylation in the CHH context (mCHH) was nearly absent, but in *ddm1*, it was only partially reduced (Figure 1). The residual mCHH could be explained by continued activity of ZMET2 and ZMET5, as the mCCH subcategory of mCHH, which is more strictly dependent on RdDM (Gouil and Baulcombe, 2016), was strongly reduced (Figure 1). These results in maize are consistent with prior analyses of double mutant of DDM1-type nucleosome remodelers in rice, which resulted in greater than 50% reduction in mCHH near genes (Tan et al., 2016). Relations between DDM1 and RdDM in Arabidopsis, however appear to be different than in maize or rice, as DDM1 is thought to have minimal if any impact on canonical RdDM in Arabidopsis (Zemach et al., 2013; Lyons and Zilberman, 2017). Mutants that lack all three of the Arabidopsis or rice chromomethylases have not yet been reported. In maize, the effect of *cmt* and *ddm1* mutants was weaker in 14 DAP-endosperm than in 14-DAP embryo or than in mature endosperm, particularly for *cmt*. Transfer of wildtype maternal products directly into developing endosperm might explain these differences.

Loss of 24nt siRNAs in mCHH islands and TIR DNA transposons agreed well with loss of methylation, both by tissue and genotype (Figure 2A and 2B). The gain of siRNAs in heterochromatin, particularly in retrotransposons, is consistent with *ddm1* mutants in Arabidopsis (Creasey et al., 2014; McCue et al., 2015). The *knob180* tandem repeat in maize is associated with an extreme form of heterochromatin (Peacock et al., 1981). *knob180* siRNAs increased up to 18-fold in abundance in *ddm1* but not in *cmt* (Figure 2B). Even though 24nt siRNAs also increased, mCHH decreased, indicating that these siRNAs did not lead to productive RdDM. The fact that production of siRNAs requires transcription indicates transcriptional derepression in *ddm1* and suggests that DDM1-type nucleosome remodelers can have roles in transcriptional silencing independent of chromomethylation.

24nt siRNAs were not completely lost in *ddm1* or *cmt.* In fact, the total number relative to miRNAs was only decreased to about half of wildtype levels (Figure 2B), and they were retained at high levels at discrete loci throughout the genome (Figure 3A). Loci that retained or gained 24nt siRNAs in *ddm1* or *cmt* embryos tended to have abundant 21 and 22nt siRNAs, even in wildtype (Figure 3B).In Arabidopsis, the additional siRNAs produced in *ddm1* mutants can direct mCHH using alternative forms of RdDM, depending on the cell type (Nuthikattu et al., 2013; McCue et al., 2015; Panda et al., 2016). Alternative forms of RdDM undoubtedly exist in maize too, but the loci that gained or retained 24nt siRNAs in *ddm1* and *cmt* embryos, and which were rich in 21 and 22nt siRNAs, had low mCHH in wildtype and even lower in mutants. This was true even for novel *ddm1* 24nt siRNA loci that did not qualify as 24nt siRNA loci in wildtype (Figure 3C).

While de novo RdDM is required for establishment of chromomethylation, once RdDM is established, the roles are reversed (Cuerda-Gil and Slotkin, 2016). Chromomethylation is maintained by CMT and not dependent on canonical RdDM. Canonical RdDM, in contrast, depends on CMT to be fully effective, likely because CMT promotes H3K9me2. The H3K9me2 binding protein SAWADEE HOMEODOMAIN HOMOLOGUE 1 (SHH1) promotes Pol IV (Law et al., 2013; Zhang et al., 2013). The fact that maize SHH1 interacts with both the Pol IV and Pol V protein complexes might make RdDM even more dependent upon H3K9me2 in maize than in Arabidopsis, where SHH1 interacts only with the Pol IV complex (Haag et al., 2014). Likewise, interactions between Pol V cofactors and DNA methylation could explain some loss of RdDM in *cmt*. The principle reason, however, for loss of RdDM may be simple dilution. The loss of methylation from heterochromatin could result in the spreading of at least one critical RdDM component from mCHH islands into the newly accessible heterochromatin (Figure 4). Rather than an increase in RdDM at new sites across the genome, we might expect a global decrease because no specific loci would reproducibly recruit the full complement of RdDM components needed to sustain DRM activity. Three lines of evidence support the dilution hypothesis. First is the greater than 50% loss of RdDM that occurs in single mutants of either *zmet2* or *zmet5* in leaf tissues (Supplemental Figure S4) (Li et al., 2015a). A mild increase in heterochromatin accessibility genomewide could have a large effect in diluting RdDM components from mCHH islands and exacerbate these single mutant phenotypes. The second is the loss of RdDM without the loss of mCHG in mCHH islands in *ddm1* mutants (Figure 1). This observation rules out the simple scenario that DDM1 is required for chromomethylation (which is required for RdDM) and suggests an alternative explanation such as the RdDM dilution model. Finally, DDM1 is not known to function directly in RdDM in Arabidopsis, while an increase in heterochromatin accessibility genomewide in *ddm1* is strongly consistent with its known function and mutant phenotypes (Gendrel et al., 2002; Zemach et al., 2013; Creasey et al., 2014; McCue et al., 2015; Lyons and Zilberman, 2017). The stronger effect of *ddm1* and *cmt* mutants on RdDM in maize could be explained by its larger genome size, as dilution of RdDM components would be less of a problem in plants like Arabidopsis with a genome one-nineteenth the size of maize.

**Figure 4:**
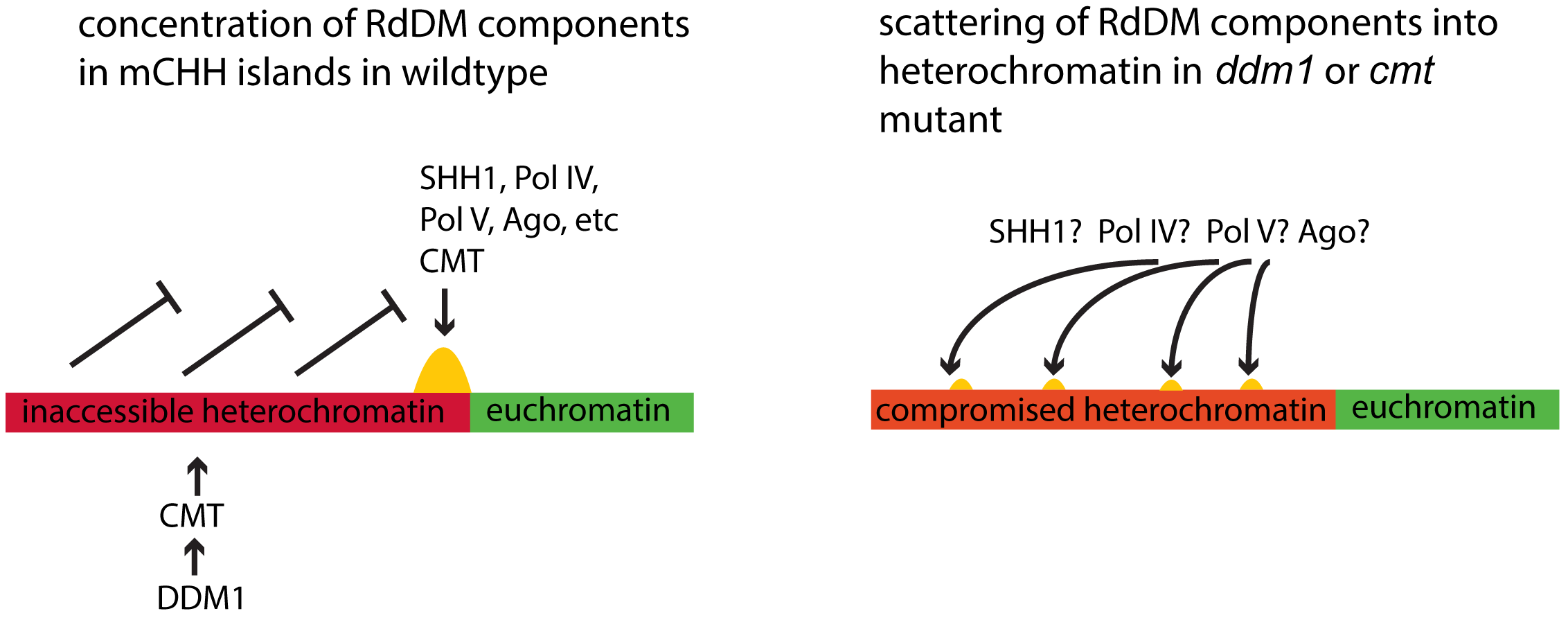
A hypothetical explanation of loss of RdDM in *ddml* and *cmt* mutants In wildtype conditions, heterochromatin is maintained in an inaccessible state that excludes RdDM. All the components required for RdDM are then concentrated at a small set of loci in the genome (mCHH islands) where they function in concert to methylate DNA. In the absence of chromomethylases or DDMl-type nucleosome remodelers, heterochromatin no longer excludes RdDM, and RdDM components are scattered over a vast area of repetitive DNA. This hypothesis would explain the absence of mCHH in the genome if even a single critical component were present at too low a concentration for productive RdDM.

## ACKNOWLEDGEMENTS

We thank Sean Trostel for assistance with genotyping, the Maize Genetics Cooperation Stock Center for providing UniformMu stocks, and Nathan Springer for providing other stocks and for comments on the manuscript. This study was supported in part by resources and technical expertise from the Georgia Advanced Computing Resource Center, a partnership between the University of Georgia’s Office of the Vice President for Research and Office of the Vice President for Information Technology. Funding for this study was provided by the National Science Foundation, #0922703 and #1118550 to RKD, and #1547760 to JIG.

## AUTHOR CONTRIBUTIONS

FFF designed and performed research. RKD designed research and wrote the paper. JIG designed and performed research, analyzed data, and wrote the paper.

## METHODS

PCR primers, alleles and gene names for all mutants used in this study are listed in Table 1. WT controls were homozygous wildtype for all four genes and were progeny of homozygous wildtype parents. Developing endosperm and embryos were collected 14 days after pollination and flash frozen in liquid nitrogen for later nucleic acid extraction. Prior to freezing, pericarps were removed from kernels and the embryos separated from the endosperm. Each endosperm was cut into two halves, one for DNA extraction and one for RNA extraction. For mature endosperm, dry kernels were soaked in 6% NaOH in water at 57°C for 8 minutes and the pericarps removed with forceps. Each mature endosperm was ground to a powder with a mortar and pestle. Frozen 14-DAP endosperms and embryos were ground with micropestles in 2mL microcentrifuge tubes without thawing. DNA was extracted from all three tissue types, each individual sample separately, with the DNeasy Plant Mini Kit (QIAGEN #69104). WGBS libraries were prepared using the methylC-seq method (Urich et al., 2015) with no more than 7 cycles of PCR amplification for endosperm and no more than 10 for embryo. For the mature endosperm libraries of Figure 1C, DNA from three individuals was combined. For all other libraries, separate libraries were made from each individual embryo or endosperm. All results shown are the average of two or three individuals per genotype, except *Zmet2/zmet2 zmet5/zmet5* in Figure S2, which is derived from a single embryo.

**Table 1:**
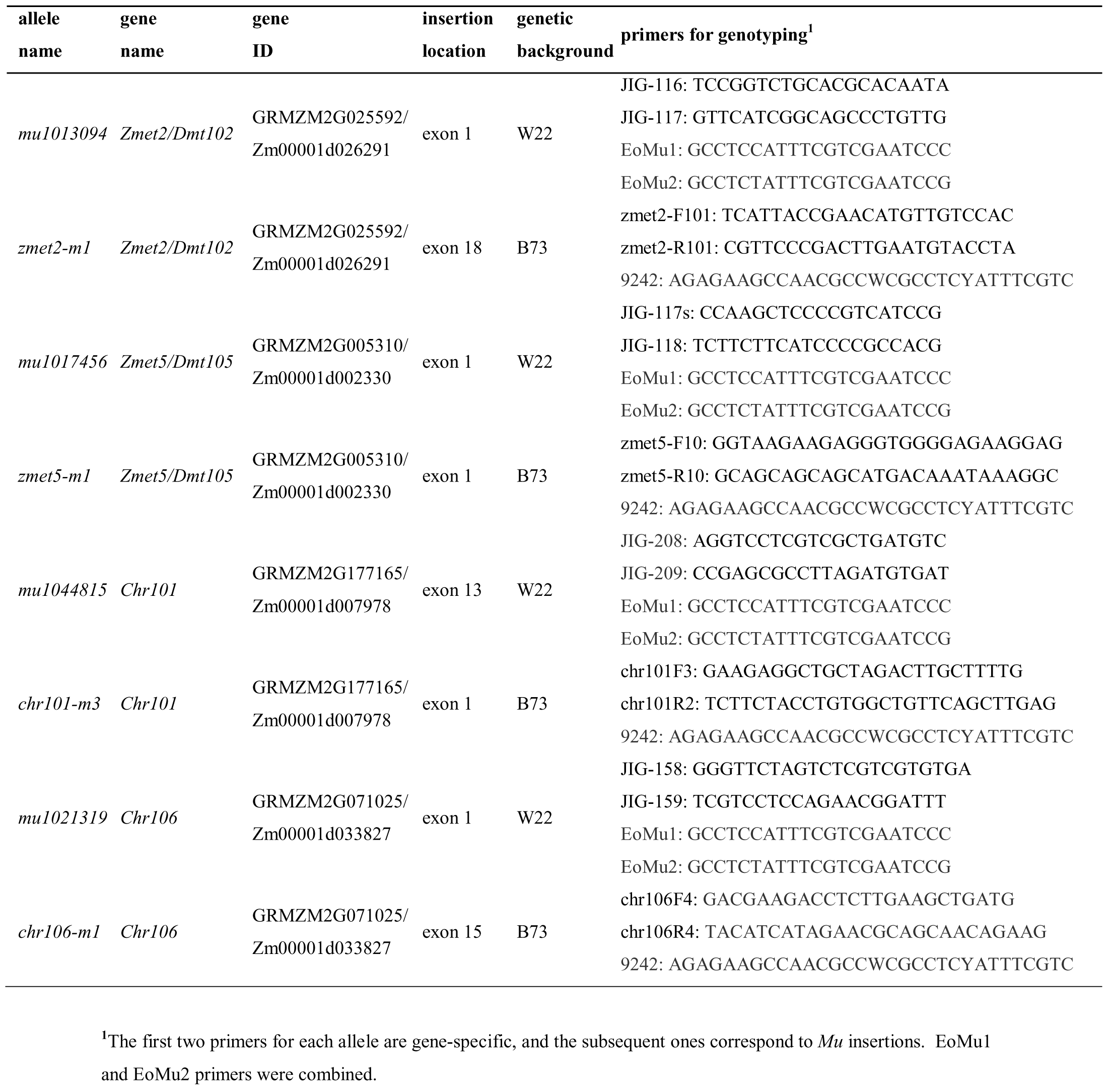
mutant alleles and genotyping primers

RNA was extracted from individual 14-DAP embryos and 14-DAP endosperm using mirVana^TM^ miRNA Isolation Kits (ThermoFisher Scientific #AM1560) using the total RNA method. For the 14-DAP endosperm, Plant RNA Isolation Aid (ThermoFisher Scientific #AM9690) was added at the lysis step. Small RNA sequencing libraries were prepared from individual embryos and endosperm (two or three for each genotype) using the NEXTflex^TM^259 cl:1 Small RNA-Seq Kit v3 (Bioo Scientific #5132-05) with 13 cycles of PCR amplification for endosperm and 17 cycles for embryo. 150nt single-end Illumina sequencing was performed on an Illumina NextSeq 500 system at the Georgia Genomics Facility, University of Georgia, Athens, GA, USA.

BS-seq reads were trimmed and quality filtered using cutadapt (version 1.9.dev1 with Python 2.7.8) (Martin, 2011), command line parameters “-q 20 ‐a AGATCGGAAGAGC ‐e .1 ‐O 1 ‐m 50”. Trimmed reads were aligned to the maize W22 references genome [citation] using BS-Seeker2 (v2.1.1 with Python 2.7.8 and Bowtie2 2.2.9) with default parameters except –m 1 to allow for a single mismatch. All libraries were aligned to the *Zea* consensus sequences of the 156bp tandem repeat *CentC* and the 180bp tandem repeat *knob180* (Gent et al., 2017) using BS-seeker2 in the same way, except up to four mismatches were allowed per read.

Small RNA-seq reads were trimmed and quality filtered using cutadapt (version 1.14 with Python 2.7.8) (Martin, 2011), command line parameters “-u 4 ‐q 20 ‐a TGGAATTCTCGGGTGCCAAGG ‐e .05 ‐O 20 ‐‐discard-untrimmed ‐m 24 ‐M 29” followed by a second trim with just “-u ‐4”. In this way adapter sequences and the four random nucleotides at each end of each RNA were trimmed and all reads outside the range of 20-25nt were removed. NCBI BLAST (version 2.2.26) was used to identify reads corresponding to the set of maize mature miRNA sequences from miRBase (Version 20, (Kozomara and Griffiths-Jones, 2011)). The blastall Expectation value was set to 1e-5. Reads corresponding to the tandem repeats *CentC* and *knob180* (Gent et al., 2017) were identified similarly, except the blastall Expectation value was set to 1e-6. The consensus sequence for each was turned into a dimer to allow reads that spanned the junctions between monomers to be identified. After removing all identified miRNAs from the small RNA reads, the remaining 20-25nt reads were mapped to the W22 genome [citation] using the BWA-backtrack (version 0.7.15 (Li and Durbin, 2009)), command line parameters “aln ‐t 8 ‐l 10”. All mapping reads were included in the set of siRNAs, including non-uniquely mapping reads. All results shown are averages from two or three individual embryos or endosperms. Whole genome coverage was calculated on 500Kb intervals and visualized using the Integrative Genomics Viewer (Thorvaldsdóttir et al., 2013).

All sequencing reads have been deposited in the Sequence Read Archive (SRA), SRP127627. Read counts for each experiment and SRA accession numbers are listed in Supplementary Tables 4 and 5.

